# Transformer-based tool recommendation system in Galaxy

**DOI:** 10.1101/2022.12.16.520746

**Authors:** Anup Kumar, Björn Grüning, Rolf Backofen

## Abstract

Galaxy is a web-based open-source platform for scientific analyses. Researchers use thousands of high-quality tools and workflows for their respective analyses. Tool recommender system predicts a collection of tools that can be used to extend an analysis. In this work, a tool recommender system is developed by training a Transformer-based neural network on workflows available on Galaxy Europe. Compared to the existing tool recommender system on Galaxy Europe that trains a recurrent neural network, the transformer-based neural network achieves two times faster convergence, has a four times lower model usage (model loading + prediction) time and shows a better generalisation that goes beyond training workflows. The scripts to create the recommendation model are available under MIT licence at https://github.com/anuprulez/galaxy_tool_recommendation_transformers.

## 1: Findings

### A. Background

A rapid increase in the number of scientific tools that achieve a variety of tasks in different fields of life sciences makes constructing workflows using these tools more complicated. Assembling such scientific tools into a workflow poses a significant challenge as the analysis represented by the workflow should incorporate scientifically significant steps and produce reproducible results. To simplify creating workflows, a tool recommender (1) in Galaxy (2) was created using good-quality workflows stored in Galaxy Europe. This recommender system trains a recurrent neural network (RNN) on existing good-quality workflows and creates a model that predicts scientific tools at each step of creating workflows. At each step, it takes into account the sequence of scientific tools or already created workflow to recommend tools. RNNs have been popular for modelling sequential data but there are a few drawbacks to such an architecture. First, the training and convergence times are significantly higher compared to Transformers (3) as it is hard to parallelise mathematical computations because of recurrent connections and large memory consumption that forces training to work in small batches of workflows. As data grows with time, it becomes more challenging to train such an architecture. Second, the trained RNN model has a larger model usage time than the Transformer. Third, it is challenging to model longer sequences and lastly, the generalisation ability of the Transformer is better compared to the RNN. To harness these benefits, the recommender model is created by training a Transformer on workflows in Galaxy Europe.

## 2: Transformers

Transformers have been used successfully in several studies to model sequential data to achieve start-of-the-art outcomes. Bidirectional Encoder Representations from Transformers (BERT) has been vastly used for modelling languages and achieves exceptional results on eleven natural language tasks (4). DNABERT creates a novel model taking cues from BERT architecture by training on DNA sequences. It is used for many downstream tasks such as the prediction of regulatory elements, promoters, splice sites and transcription factor binding sites with high accuracy (5). ProteinBERT improves the BERT architecture to model protein sequences and achieves excellent results on various tasks such as predicting protein functions and Gene Ontology (GO) annotations (6). Transformer makes use of the attention mechanism (3) to learn representations of sequential data. Each token in the sequential data is assigned a weight that is used collectively to compute predictions. These weights are represented by real numbers. A token with a larger magnitude of weight is more important than one with a smaller magnitude in prediction tasks. The architecture of a transformer has several components. Two broad components are the encoder and decoder (3). For sequential data that consists of input and output sequences, the encoder learns the representation of input sequences and the decoder jointly trains on the learned representation of input sequences and the output sequences. In our work, only the encoder part of the Transformer is used to learn a representation of sequences of tools and recommend tools for tools or tool sequences. The architecture of the Transformer used for creating the Galaxy tool recommender is discussed in the Architecture section (Figure 1).

**Fig. 1.**
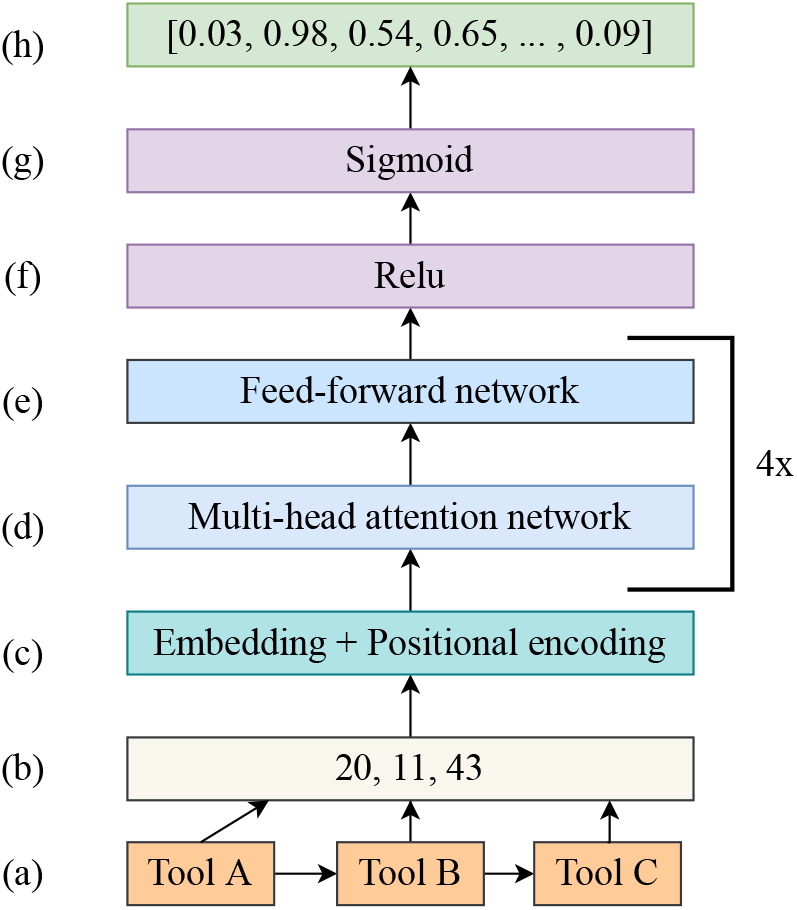
Neural network architecture of Transformer used for recommending Galaxy tools. Figures 1a and 1b represent how a sequence of tools is transformed into a sequence of integers. Figures 1c - 1g represent several different neural network layers through which sequences of tools are passed to learn mapping with output tools. Figure 1h represents the output in the form of a real-valued vector.

### A. Architecture

The encoder part of the Transformer consists of multiple layers as shown in Figure 1. The first ones are embedding and positional encoding layers (Figure 1c). These two layers work together to encode tools, where each tool is represented by a unique integer (Figure 1b), in workflows (Figure 1a). Tools are tokens that are sequentially arranged in workflows. The encodings of tools contain real-valued vectors that conserve the relative positions of tools in workflows. These encodings of sequences of tools are passed to a neural network consisting of multi-head attention and feed-forward layers (Figures 1d and 1e). The term 4x in Figures 1d and 1e represent 4 attention networks working together in parallel. Collectively, they compute representations of sequences based on attention weights. In the last step, these representations are passed to a feed-forward neural network for the prediction task (Figures 1f and 1g). In our work, the prediction task is multi-label, multi-class classification as each sequence of tools may be extended with one or more tools. The predicted output is a vector of sigmoid scores (Figure 1h) that are used for recommending top N tools. The higher the sigmoid score, the higher the probability of a tool being correctly recommended.

## 3: Implementation

The architecture of the Transformer used in our work is implemented in TensorFlow (7) using a functional API. The neural network architecture of the Transformer is trained on a virtual machine with an Nvidia GPU for 35,000 iterations with a batch size of 128. Adam optimiser (8) is used for minimising the binary cross-entropy (9) error with a standard learning rate of 0.001. The output layer of the feed-forward network has a sigmoid activation. All the sequences (approximately 500,000) of tools are divided into train and test sets. 80% of the sequences are used for training (approximately 400,000) and the rest for testing (approximately 100,000) the trained model. The codebase (10) is written in Python 3.9. Weight matrices of the trained model are stored as an H5 file. These weights along with Transformer architecture are used for reconstructing the model for recommending tools on Galaxy Europe. An API is used for recommending tools (11). It is possible to create the Transformer model on Galaxy Europe by running a Galaxy tool (12) that executes the same scripts as in (10). A script (13) is used to collect workflows from any Galaxy server.

## 4: Results

### A. Transformer convergence time

Transformer and RNN architectures [ref from old paper] are trained on the same dataset and their precision@k score collected over 35,000 training iterations on the test data are compared in Figure 2. Precision@k is used as the prediction metric and is popular for evaluating recommender systems (14–16). The precision@k scores are averaged over 5 experiment runs. In each run, the train and test datasets are randomly created but have the same size. In Figure 2, the precision@k score is shown in green for the Transformer and in red for the RNN. The training iterations in both experiments are set to 35,000. Figure 2 shows that the Transformer architecture converges two times faster than the RNN. The Transformer converges to the precision@k score of approximately 0.98 already at iteration 10,000 while the RNN achieves similar precision@k only at iteration 20,000. All other parameters of the experiment remain the same.

**Fig. 2.**
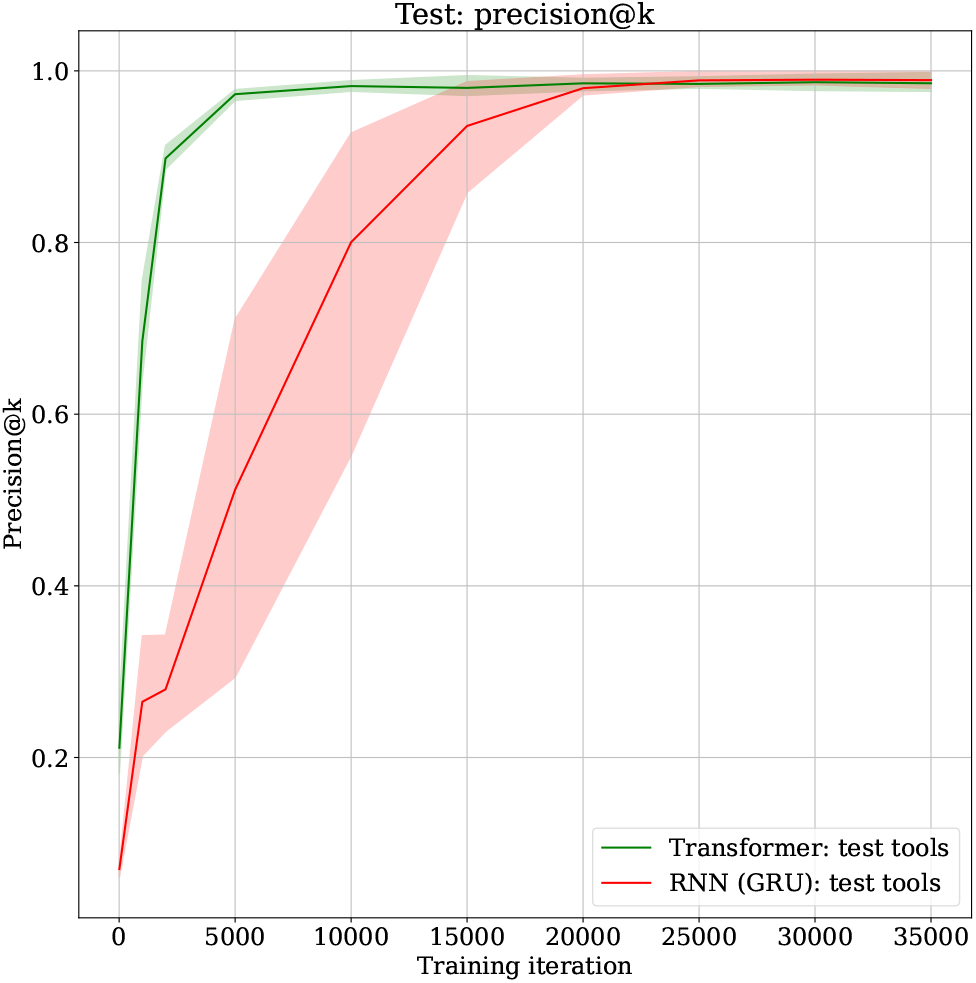
The figure shows how the precision@k metric for evaluating a recommender system improves over training iterations for the Transformer (shown in green) and RNN (shown in red) architectures. The precision@k values are averaged over 5 experiment runs each time using a random set of training and test sequences of tools. The shaded regions show standard deviation of both models.

### B. Transformer usage time

Transformers and RNN architectures are compared for their respective model loading and prediction time measured over 35,000 training iterations as shown in Figure 3. The model usage time (= model loading + prediction) of the Transformer is less than 0.5 seconds for a sequence of tools while for the RNN it is approximately 2 seconds. The model usage time is averaged over all the sequences of tools in the test dataset. It can be concluded that the Transformer’s prediction is approximately 4 times as fast as the RNN.

**Fig. 3.**
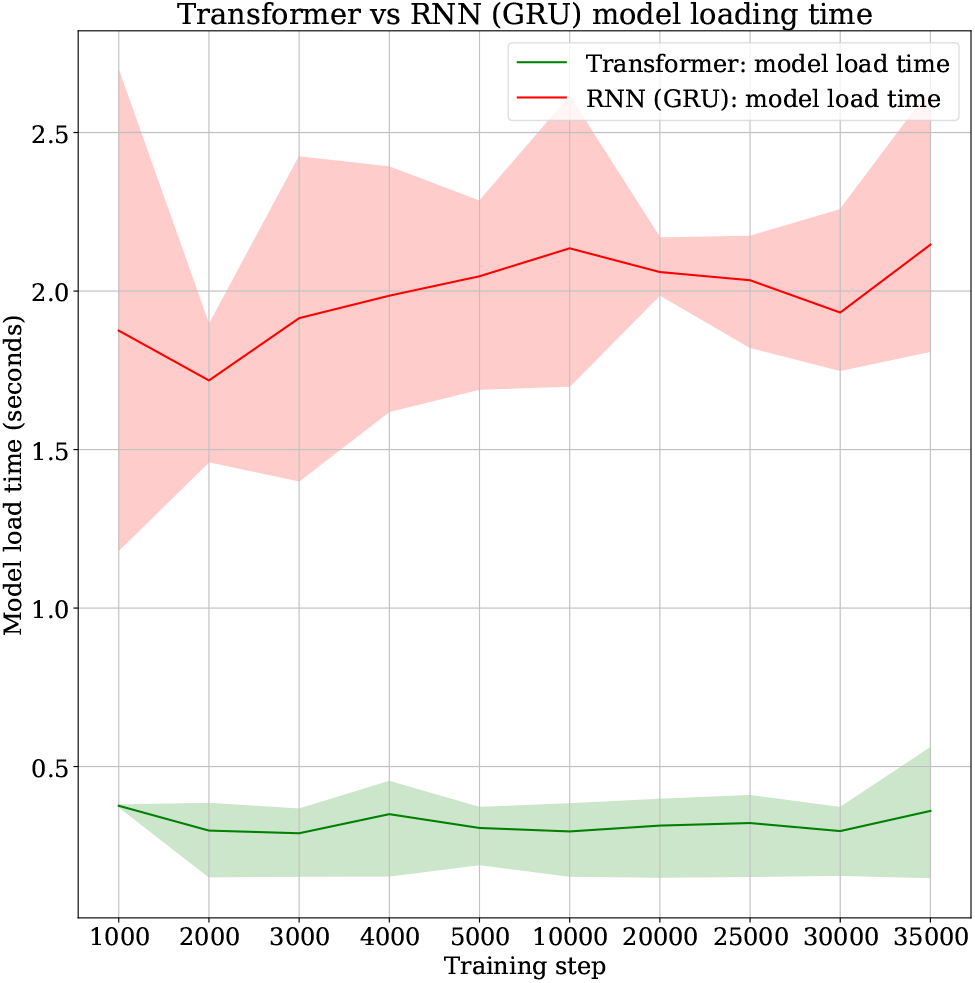
The figure shows a comparison of the model usage (model loading + prediction) time of the Transformer (shown in green) and RNN (shown in red). The time is averaged over 5 experiment runs and the shaded region show standard deviation.

### C. Generalisation beyond training data

Trained models of the Transformer and the RNN have been used to recommend the top 20 tools using tools and sequences of tools belonging to multiple scientific analyses as shown in Table 1. The ground truth column shows true tools, extracted from training workflows, for a tool or a tool sequence in the second column. The third and fourth columns show the recommended tools, predicted by the Transformer and RNN models, respectively. The tools that are shown in bold are those recommended tools that are compatible with the respective tool or tool sequence, in the second column, but such connections are not available in the training workflows. To elaborate, “freebayes” (18) tool (shown in the first row of Table 1) in the third column can be used on the output datasets produced by “snpeff_sars_cov_2” (17) tool to extend a workflow for variant calling analysis. But this connection does not exist in the training workflows used to train the Transformer model as it is absent from the ground truth recommendation of the “snpeff_sars_cov_2” tool. Similarly, tools such as “mimodd_varcall” (20), “snpfreqplot” (19), “gemini_load” (28) and “vcfcombine” (21) can be used to extend a scientific analysis after using “snpeff_sars_cov_2” tool but its connections to these recommended tools do not exist in the training workflows. RNN model also recommends 3 tools that are beyond the seen training workflows but the Transformer recommends 5 such tools. Another example is from the Proteomics field where the ground truth recommendations are correct by both models but Transformer recommends many other tools that are from the Proteomics field and can be used to extend the “cardinal_preprocessing → cardinal_segmentations → mass_spectrometry_imaging_filtering” (27) (fourth row of Table 1). Similar instances of better generalisation of the Transformer are found for other scientific analyses such as Single-cell and Deep learning where the sets of high-quality recommendations by the Transformer that go beyond training workflows are significantly larger than that by the RNN while the ground truth recommendations by both the models remain the same. Therefore, from Table 1, it can be concluded that the Transformer model generalises better than the RNN model in recommending tools as it predicts those tools that are valid but have not been seen while training.

**Table 1.**
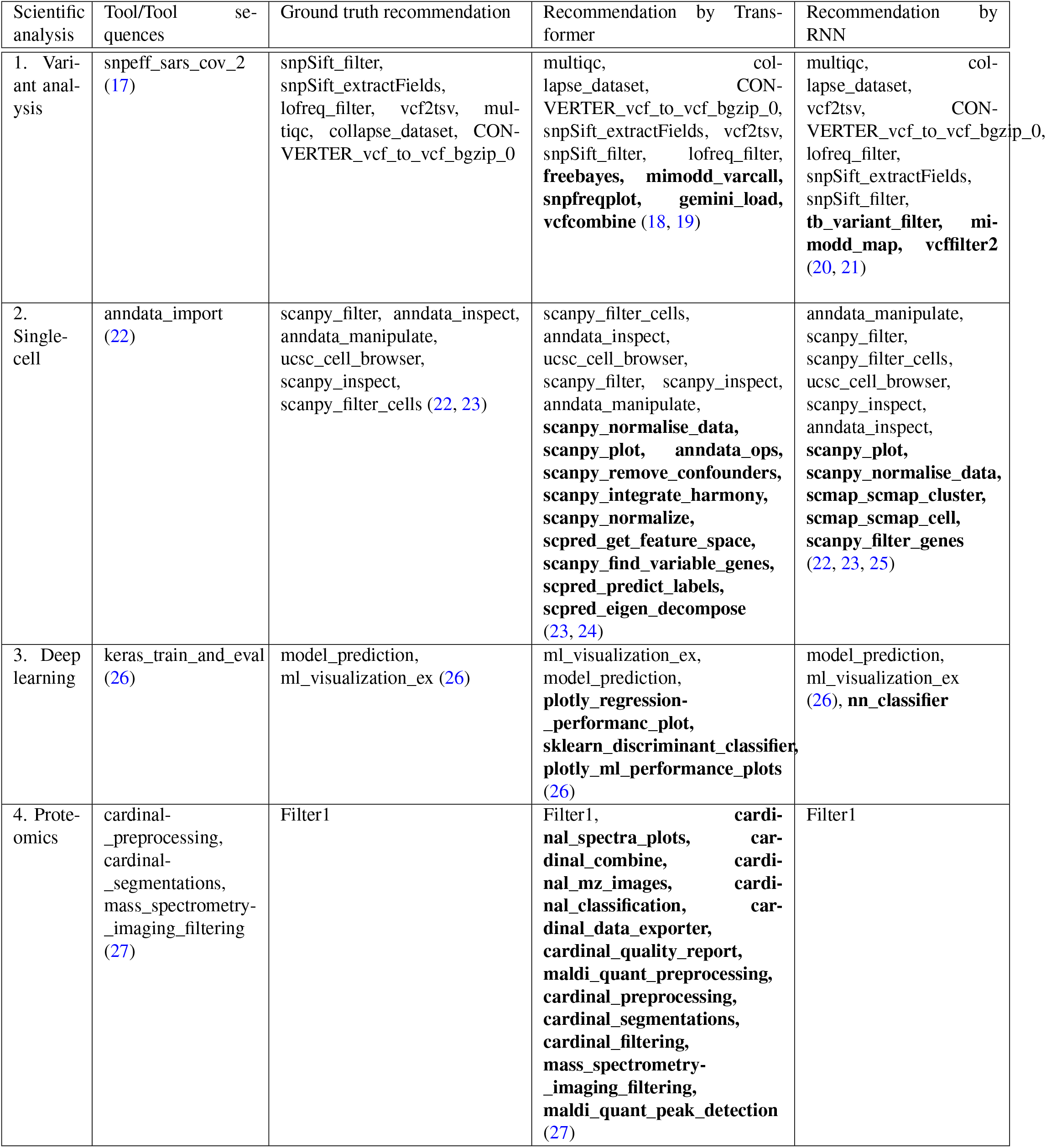
Comparison of recommendations by Transformers and RNN

## 5: Discussion and Summary

Transformer neural network outperforms RNN on multiple key parameters such as 2 times faster in model convergence, 4 times faster in prediction and recommending high-quality tools that are beyond training workflows. Being faster in model convergence enables faster creation of a new model and the faster prediction helps in showing recommendations instantly on Galaxy Europe. Being better parallelisable allows the Transformer model to integrate a larger volume of training data in future. Motivated by these benefits in this work, we propose to replace the RNN model with a Transformer model for recommending tools on Galaxy Europe. To create a new recommendation model, workflows should be extracted from a Galaxy server using the script in (13) and then a new model can be created using one of these ways, either by running a Galaxy tool or by running the codebase (10) locally or on a remote computer having memory of at least 20 GB.

## 6: Availability of supporting source code and requirements

Project name: Transformed-based tool recommendation system in Galaxy

Project home page: https://github.com/anuprulez/galaxy_tool_recommendation_transformers

Operating system: Linux

Programming languages: Python, XML

Licence: MIT License

## 7: Declarations

## A. List of abbreviations

GB: Gigabyte
AI: Artificial intelligence
RNN: Recurrent neural network
GPU: Graphical processing unit
BERT: Bidirectional Encoder Representations from Transformers

## C. Author Approvals

All authors have seen and approved the manuscript, and it has not been accepted or published elsewhere.

## D. Authors’ contributions

Authors’ contributions follow the order of names.

## E. Acknowledgements

We thank the Galaxy community, especially Galaxy Europe for their support.

